# Beneficial early, harmful later: age-dependent effects of testicular factors on male mortality

**DOI:** 10.64898/2026.10.19.760267

**Authors:** Laurence T. Maeyens, Nisi Jiang, Ziying Xu, Vivian Diaz, Randy Strong, Shangang Zhao, James F. Nelson

## Abstract

Men live shorter lives than women across nearly all human populations, a disadvantage that extends across most biological causes of death and that cannot be fully explained by behavioral or environmental differences. This suggests that fundamental biological mechanisms contribute to elevated male mortality. Our previous work implicated testicular factors as a major contributor, showing that prepubertal castration extends male lifespan to that of females in UM-HET3 mice, a genetically heterogeneous stock. However, it remains unknown when testicular factors begin to exert their detrimental effects on mortality. Here, we castrated UM-HET3 males before puberty (20 days), after puberty (60 days), or immediately before the typical rise in male mortality (240 days). Castration at all three ages prevented the early male mortality and increased median lifespan by 19.3%, 25.8%, and 32.9%, respectively. The comparable survival benefits of prepubertal and postpubertal castration indicate that exposure to the pubertal surge in testicular factors, including male sex hormones, is not responsible for establishing the male longevity disadvantage. Remarkably, castration at 240 days not only preserved the survival benefit of earlier castration but produced significantly greater survival than earlier castration, suggesting that testicular exposure before 240 days is actually beneficial. The immediate prevention of excess male mortality following castration at 240 days also indicates that the responsible testicular factors act rapidly and that their continued presence is necessary to sustain elevated male mortality. Longitudinal body-composition measurements further revealed that castration shifted males toward a female-like phenotype, characterized by reduced lean mass and continued fat accumulation into later life. Our findings reveal opposing age-dependent effects of testicular factors on survival and narrow the potential therapeutic window to later adulthood.

## Introduction

Men live shorter lives than women across virtually all human populations, with a particularly pronounced male mortality disadvantage during early-to-mid adulthood^1;2^. Although men experience greater mortality from external causes, including accidents, homicide, and suicide, they explain only approximately one-third of the sex difference ^3^. Indeed, the male mortality disadvantage extends across most major age-related diseases, including cardiovascular disease and cancer ^3^. Importantly, the male mortality disadvantage persists and becomes even more pronounced after adjustment for 17 demographic, socioeconomic, behavioral, and health-related factors, including smoking ^4^. Together, these findings suggest that fundamental biological sex differences contribute to the male longevity disadvantage beyond differences in behavior and environmental exposures.

Our previous work identified testicular factors as an important contributor: using genetically heterogeneous UM-HET3 mice, which recapitulate human patterns of sex-specific survival ^5^, removal of these factors by castration extended male lifespan to match that of females^6^. This longevity benefit appears to be evolutionarily conserved, with a recent large-scale analysis associating castration with increased male survival across 117 mammalian species^7^, and historical human studies suggesting similar benefits^8;9^. However, how testicular factors ultimately promote aging and reduce lifespan remains unclear. Must castration occur before puberty to fully eliminate these sex differences in longevity? If not, how late in life can castration occur and still confer the same survival benefit? These questions have important translational implications. If early-life exposure to testicular factors irreversibly establishes the heightened mortality risk of males, then interventions initiated later in life may be ineffective. Furthermore, the development of castration mimetics, i.e. interventions that selectively block the lifespan-limiting effects of testicular factors while preserving reproductive function, might require administration early in life, potentially even before puberty. Such a narrow therapeutic window would limit the feasibility and translational potential of this approach in humans.

There are several potential ways in which exposure to testicular factors could have long-term effects on male health and survival. First, the pubertal surge in sex hormones is known to exert lasting organizational effects on behavior and physiology^10^, raising the possibility that exposure to testicular hormones during puberty establishes persistent changes that ultimately shorten male lifespan. Second, testicular factor exposure may promote damage beginning early in life, with these effects accumulating over time and eventually contributing to the elevated mortality risk observed in males.

To test these possibilities, we castrated genetically heterogeneous UM-HET3 males at different ages to determine how the duration and timing of testicular factor exposure influence health and longevity. Prepubertal castration at day 20 (ORX20), as in our previous study^6^, served as a positive control alongside a female control group. Postpubertal castration at day 60 (ORX60) allowed us to test whether exposure to testicular factors during puberty is necessary for their long-term detrimental effects, whereas castration at day 240 (ORX240), right before the typical rise in male mortality risk, tested whether castration could still prevent excess male mortality despite prolonged prior exposure, and whether this mortality disadvantage depends on the ongoing presence of testicular factors. Each of the three castration groups was accompanied by a littermate-matched sham-operated control group.

## Results

### Male lifespan is not limited by pubertal testicular factor exposure

Castration at all three ages prolonged survival, with particularly pronounced effects on early mortality (Figure 1A-C). Because the survival curves indicated that the effect of castration varied over time, we tested the proportional hazards assumption and found evidence of non-proportional hazards for all castration groups. We therefore complemented the log-rank test with the Peto–Peto modification of the Gehan–Wilcoxon test, which places greater weight on earlier survival differences and was better suited to the early separation observed between the survival curves^11^.

**Figure 1.**
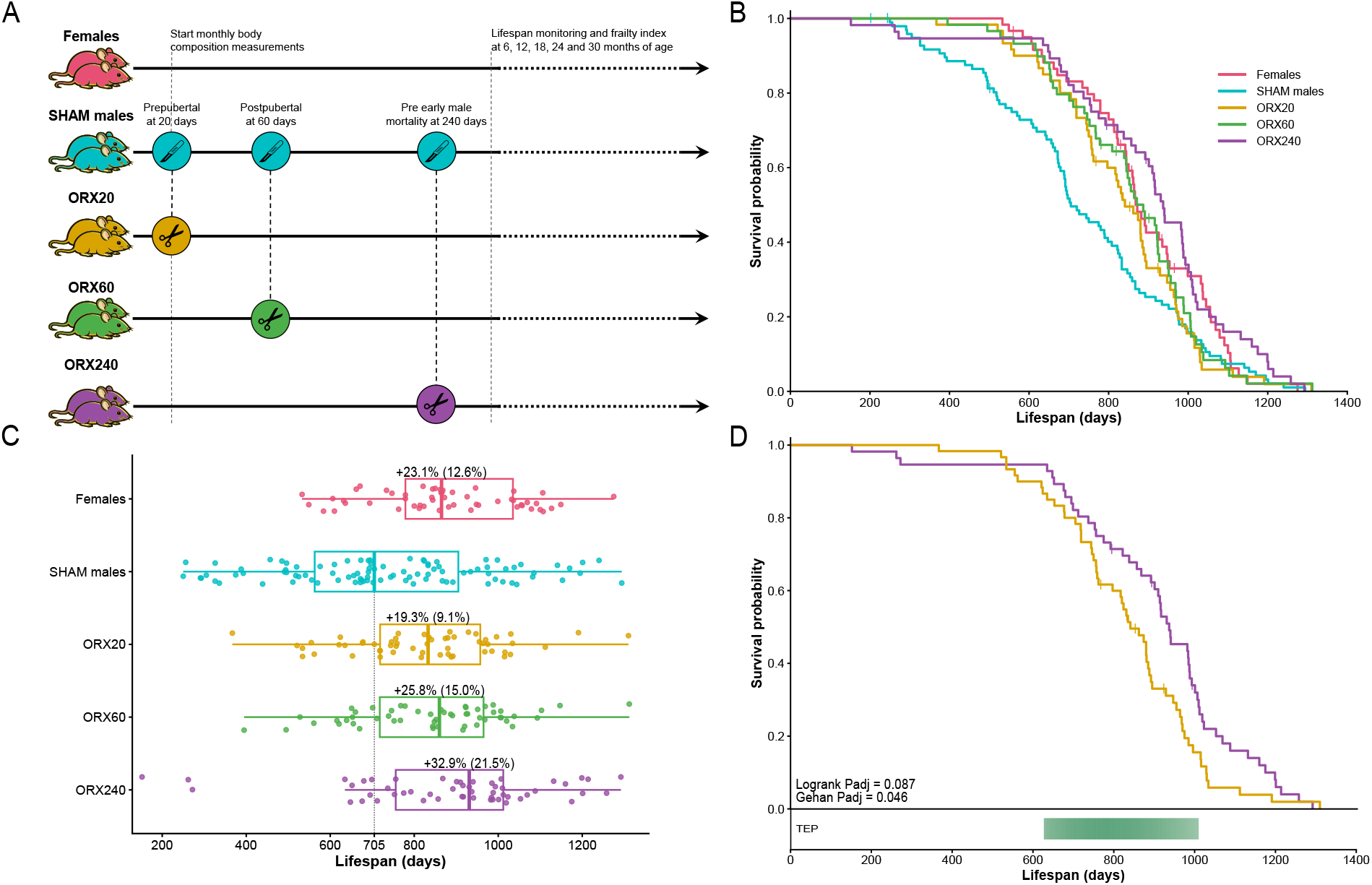
Castration at several ages extends male lifespan and reveals age-dependent effects of testicular factors on survival. A) Experimental design. Genetically heterogeneous UM-HET3 males underwent castration at approximately 20 days (ORX20; prepubertal; n=60), 60 days (ORX60; postpubertal; n=59), or 240 days of age (ORX240; prior to the typical rise in male mortality; n=56). Sham-operated males (n=99) and untreated females (n=60) served as controls. B) Kaplan–Meier survival curves for females, pooled SHAM males, ORX20, ORX60, and ORX240 mice. Castration at all three ages reduced the early-to-midlife male mortality disadvantage and increased median lifespan relative to SHAM males. C) Points represent individual animals that died naturally (censored animals excluded), with boxplots showing medians and interquartile ranges. Percentages indicate changes in KM median lifespan relative to pooled SHAM males, or the historical male KM median of 771 days (in parentheses). D) Kaplan–Meier survival curves comparing ORX20 and ORX240 mice. ORX240 mice exhibited greater survival than ORX20 mice, with the Temporal Efficacy Profiler (TEP) identifying a significantly reduced mortality hazard for ORX240 between 627 and 1,010 days of age. Overall and pairwise survival differences were evaluated using log-rank and Peto–Peto modified Gehan–Wilcoxon tests, with P values adjusted using the Benjamini–Hochberg procedure; TEP results were based on the asymptotic confidence-interval method with bootstrap confirmation.

Prepubertal castration (ORX20) increased median lifespan by 19.3% relative to pooled SHAM males (841 vs. 705 days; Gehan, P = 0.0211; log-rank, P = 0.2364; Figure 1B and C), reproducing our previously observed lifespan extension toward female levels following prepubertal castration^6^. The median lifespan of the pooled SHAM cohort was lower than both that observed in our previous SHAM cohort (771 days) ^6^ and that of historical male controls from the University of Texas site of the Intervention Testing Program (ITP; 771 days) (Supplementary Figure 1A). The three SHAM groups were pooled because their survival distributions did not differ significantly (Supplementary Figure 1B). Despite the lower median lifespan, the current SHAM cohort did not differ significantly from historical controls (Gehan, P = 0.6834; log-rank, P = 0.6736), with inspection of the survival curves indicating that the difference in median lifespan was largely driven by a modest localized decrease in survival near the 50% survival threshold (Supplementary Figure 1A). The comparatively low SHAM median therefore may have magnified the percentage increase in median lifespans observed in the current cohorts. For this reason, Figure 1C additionally presents percentage changes relative to the historical male median, providing a complementary estimate of the average difference in median lifespans. This value may be more representative, as using the historical male median of 771 days as a reference, the corresponding increase for ORX20 was 9.1% (Figure 1C). Compared to 19.3%, this estimate closely matched our previous study, in which prepubertal castration increased median lifespan by 10.2% (771 versus 850 days). In that substantially larger cohort (∼220 animals per group), the survival difference also became significant by log-rank analysis.^6^

To determine whether exposure to testicular factors during puberty limits male lifespan, we also castrated mice after puberty at 60 days of age (ORX60) (Figure 1A-C). ORX60 increased median lifespan by 25.8% relative to SHAM males (887 vs. 705 days; Gehan, P = 0.0073; log-rank, P = 0.1657). Importantly, although median lifespan was numerically greater in ORX60 than in ORX20 mice, survival did not differ significantly between these groups (Gehan, P = 0.4461; log-rank, P = 0.4696). This demonstrates that exposure to male sex hormones during puberty is not responsible for the shorter lifespan of males.

### Testicular factor exposure before 240 days benefits survival, whereas continued exposure later in life is detrimental

UM-HET3 males begin to show appreciable mortality shortly after 240 days of age (Supplementary Figure 1A). We therefore tested whether castration immediately before the onset of this early male mortality could still confer a survival benefit. ORX at 240 days (ORX240) was as effective as ORX at 20 and 60 days in eliminating the early increase in male mortality. Besides eliminating early male mortality, it further extended median male lifespan with a 32.9% increase compared to SHAM (937 vs. 705 days; Gehan, P = 0.0005; log-rank, P = 0.0145), producing a significantly longer lifespan than ORX20 (Figure 1D; Gehan, P = 0.0457; log-rank, P = 0.0869) and a trend with ORX60 (Gehan, P = 0.1533; log-rank, P = 0.1749). Note that P values were BH-adjusted for 10 comparisons; unadjusted Gehan/log-rank P values were 0.0228/0.0261 (ORX240 vs. ORX20) and 0.1074/0.0875 (ORX240 vs. ORX60). This importantly suggests that testicular factor exposure before 240 days is actually beneficial to male survival, whereas exposure only after 240 days becomes detrimental. To determine when the survival advantage of ORX240 over earlier castration emerged, we applied the Temporal Efficacy Profiler (TEP) ^12^, which estimates age-specific mortality hazard ratios and identifies intervals during which the hazard ratio differs significantly from 1. Using the asymptotic method, ORX240 exhibited significantly lower mortality hazard than ORX20 between 627 and 1,010 days of age (Figure 1D). Thus, exposure to testicular factors before 240 days was associated with an additional survival benefit much later in life, up to the 90^th^ percentile of lifespan, compared to the survival benefit achieved by pre- and post-pubertal castration. This suggests that testicular factor exposure during early life or adulthood may confer lasting survival benefits that persist after castration. Together, these findings provide causal evidence that testicular factor exposure during puberty and early adulthood is not detrimental to male survival and may actually be beneficial, arguing against an “earlier is better” model for the longevity effects of castration.^7^

### Castration shifts male body composition toward a female-like phenotype

Body composition differed markedly between intact males and females across the lifespan (Figure 2, Supplementary Figures 2–4). SHAM males exhibited a characteristically lean phenotype, maintaining greater lean mass and substantially lower fat mass than females throughout most of adulthood. Castration shifted both compartments toward the female phenotype, reducing lean mass and increasing fat mass relative to SHAM males. This feminization was evident following castration at all three ages, demonstrating that testicular factors actively maintain the male-specific high-lean, low-fat body composition well beyond puberty. Fat mass showed a particularly pronounced sex-dependent trajectory. Females and castrated males continued to accumulate fat until beyond ∼60% of their lifespan, whereas SHAM males stopped accumulating fat much earlier, reaching a plateau at approximately 300 days of age, or ∼40% of lifespan (Figure 2C). Thus, the continued presence of testicular factors restricts fat accumulation during mid- and later life.

**Figure 2.**
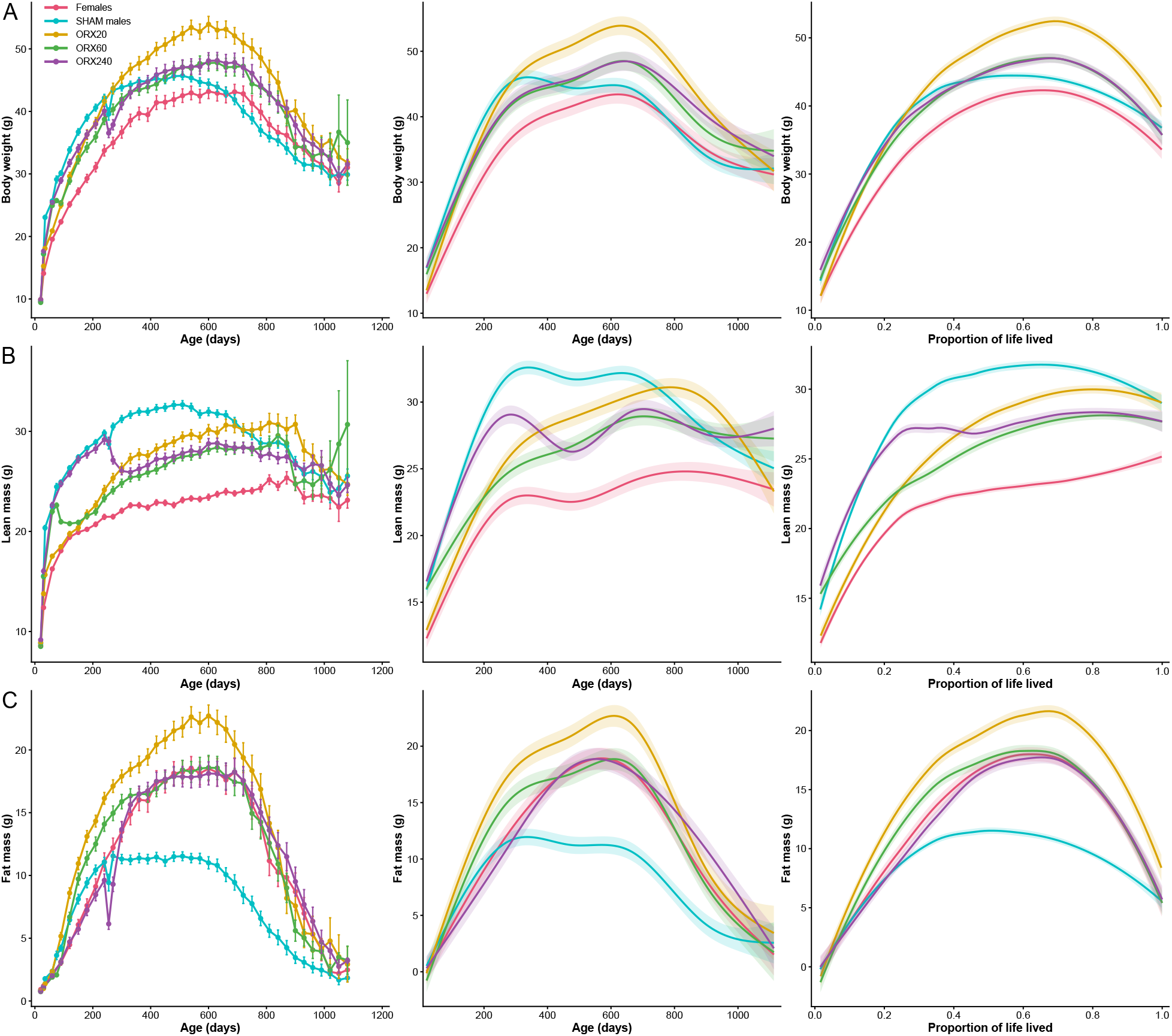
Castration shifts male body-composition trajectories toward a female-like phenotype throughout adulthood. Longitudinal trajectories of (A) body weight, (B) lean mass, and (C) fat mass in females, SHAM males, ORX20, ORX60, and ORX240 mice. Left, measurements summarized at nominal study ages and presented as group means ± s.e.m. Middle, age-dependent trajectories estimated from generalized additive mixed models (GAMMs) using the exact age at measurement values for each datapoint; solid lines represent predicted group trajectories and shaded regions represent 95% confidence intervals. Right, trajectories plotted as a function of proportion of life lived, calculated as age at measurement divided by eventual lifespan, restricted to animals that died naturally, and trajectories are shown using group-specific LOESS curves with 95% confidence intervals. Castration reduced lean mass and increased fat mass relative to SHAM males, shifting body composition toward the female phenotype. Following ORX240, body composition rapidly transitioned from an intact-male trajectory toward that of the earlier-castrated groups, demonstrating continued dependence of adult body composition on the presence of testicular factors.

The magnitude and trajectory of these changes depended on the timing of castration. Before castration, ORX240 males accumulated lean mass similarly to intact males. Following castration at 240 days, however, lean mass decreased and subsequently converged with ORX60, indicating that the amount of lean mass at any given age depends largely on the presence of testicular factors at that moment rather than earlier exposure. Surprisingly, ORX20 males maintained greater lean mass than the later-castrated groups during later life despite never experiencing pubertal testicular hormone surges. ORX20 also exhibited the greatest increase in fat mass and consequently became the heaviest group throughout much of adulthood. In contrast, ORX60 and ORX240 showed remarkably similar long-term body-composition trajectories, despite the additional 180 days of testicular hormone exposure in ORX240 animals. Together, these patterns indicate that adult body composition is highly responsive to the in situ hormonal environment rather than being irreversibly programmed by testicular factor exposure early in life.

Castration at 60 and 240 days was accompanied by an acute, transient reduction in both lean and fat mass around the time of surgery, followed by recovery and establishment of the castrated phenotype. This was particularly apparent following ORX240, where both compartments decreased sharply after surgery before recovering. A comparable decline was not readily distinguishable after ORX20, likely due to the rapid early growth period. Notably, after recovery from ORX240, its fat-mass trajectory almost completely overlapped that of females, despite substantially different body composition before surgery. This striking convergence further demonstrates the reversibility of the male body-composition phenotype and the strong dependence of adiposity on ongoing testicular factor exposure.

Finally, all groups exhibited substantial late-life body-weight loss. This decline was driven predominantly by a pronounced depletion of fat mass, whereas reductions in lean mass were comparatively modest. Thus, although castration substantially altered the amount and trajectory of fat accumulation across adulthood, late-life weight loss was characterized by loss of these adipose reserves across groups. Together, these findings demonstrate that testicular factors continuously maintain the characteristic lean, low-fat male phenotype, whereas their removal rapidly and persistently shifts body composition toward a female-like state, even when castration is delayed.

### The survival advantage of later castration is not accompanied by improved grip strength or reduced frailty

The greater lifespan of ORX240 relative to earlier-castrated males raised the possibility that testicular factor exposure during early adulthood, although detrimental when maintained later in life, may confer beneficial physiological effects before castration that contribute to subsequent survival. Body-composition trajectories provided an initial rationale for this hypothesis (Figure 2). Before castration, ORX240 males retained the characteristic intact-male phenotype, including greater lean mass, for substantially longer than animals castrated at 20 or 60 days. Thus, the additional period of testicular factor exposure experienced by ORX240 animals could potentially promote somatic or musculoskeletal development during early adulthood and establish greater physiological reserve before the testes become detrimental to survival. Such an effect would provide a potential explanation for why delaying castration until 240 days produced greater longevity than castration at 20 days (Figure 1D).

However, the later body-composition trajectories did not support a simple model in which ORX240 animals survived longer because they permanently retained more lean mass. Following castration, ORX240 lean mass declined and converged with that of ORX60, whereas ORX20 unexpectedly maintained greater lean mass than both later-castrated groups at older ages. Nevertheless, greater lean mass does not necessarily correspond to greater muscle function, and a transient period of androgen exposure during growth or early adulthood could conceivably produce lasting functional consequences even after gross body composition changes. Longitudinal assessments of forelimb grip strength, total grip strength, and frailty enabled us to test the possibility that early-adult testicular factor exposure established a functional reserve that contributed to the later survival advantage of ORX240.

Both measures of grip strength declined substantially with age in all groups, whereas frailty progressively increased (Figure 3A–C), as expected. However, ORX240 animals did not exhibit a consistent advantage in either forelimb or total grip strength relative to animals castrated earlier. Likewise, frailty did not accumulate more slowly in ORX240 animals. Aligning these phenotypes by proportion of life lived produced similar patterns: grip strength declined and frailty increased as animals approached the end of life, without evidence that ORX240 animals maintained superior physical function or delayed frailty relative to ORX20 or ORX60 animals.

**Figure 3.**
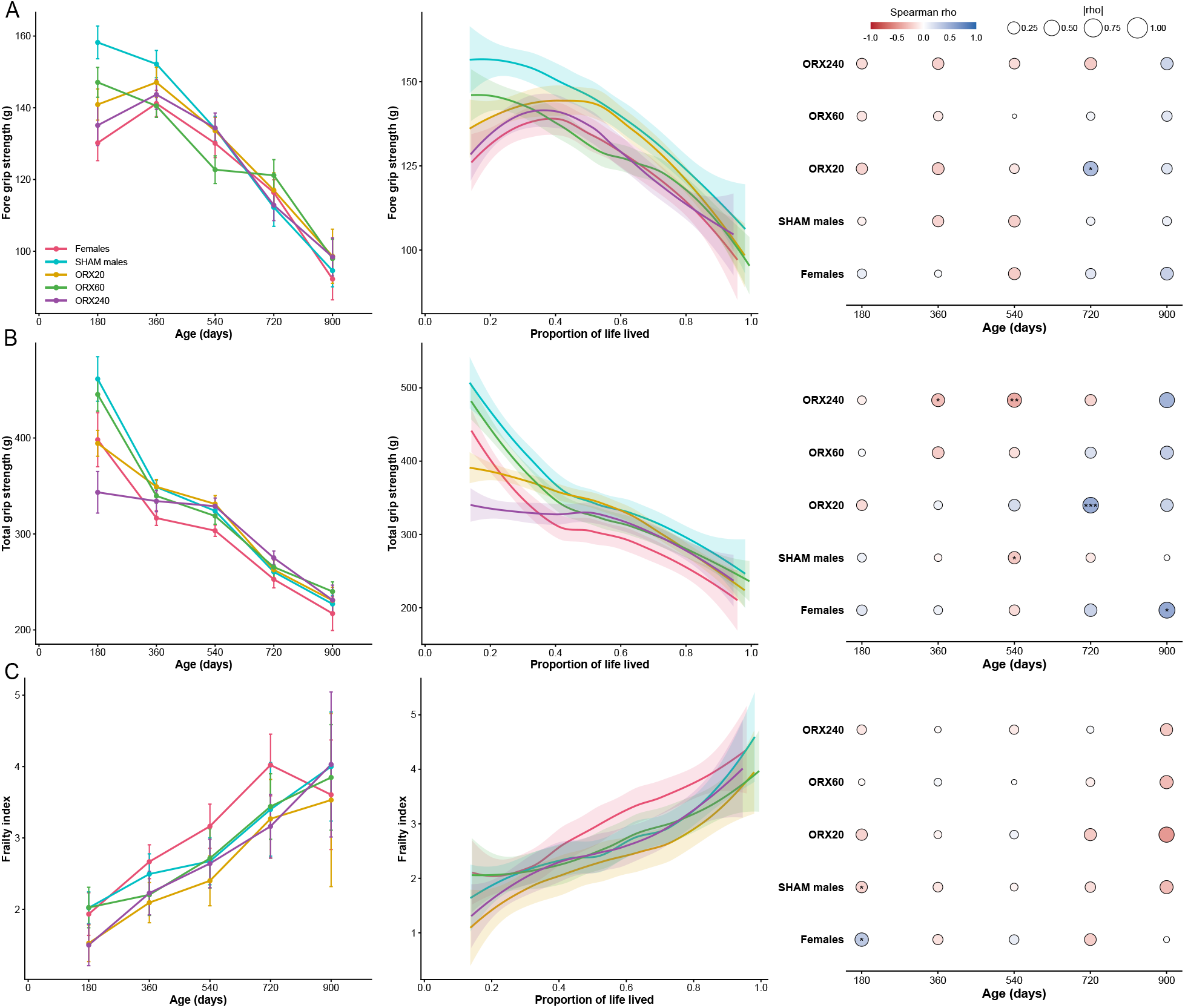
The survival advantage of later castration is not accompanied by improved grip strength or reduced frailty. Longitudinal changes in (A) forelimb grip strength, (B) total grip strength, and (C) frailty index in females, SHAM males, ORX20, ORX60, and ORX240 mice. Left, measurements across chronological age, presented as group means ± 2 s.e.m. Middle, trajectories plotted as a function of proportion of life lived among animals that died naturally; solid lines represent group-specific LOESS fits and shaded regions represent 95% confidence intervals. Right, Spearman rank correlations between each phenotype and eventual lifespan within each treatment group and measurement age. Circle size represents the magnitude of the Spearman correlation coefficient and color indicates the direction of the association. Correlation analyses were restricted to animals with known natural lifespans. Asterisks indicate P values (*P < 0.05, **P < 0.01, ***P < 0.001). Grip strength generally declined and frailty increased with advancing age, but ORX240 mice did not exhibit consistently better physical function, reduced frailty, or stronger associations between these phenotypes and lifespan than animals castrated earlier.

Notably, differences in grip strength were already apparent among treatment cohorts before all interventions had occurred. At approximately 180 days of age, ORX240 animals exhibited lower grip strength than grouped SHAM males despite both groups still having testes. Because treatment cohorts were generated sequentially rather than through contemporaneous randomized assignment, this pre-existing difference indicates that cohort-level factors independent of castration contributed to variation in physical performance. Accordingly, absolute between-group differences in grip strength, particularly those present before the relevant intervention, should be interpreted cautiously.

We therefore additionally asked whether physical function predicted eventual lifespan within each treatment group, reducing the influence of pre-existing differences between treatment cohorts. Spearman correlations between grip strength or frailty and lifespan were generally modest and varied across ages and groups (Figure 3A–C (right panels); Supplementary Figure 5). Higher grip strength was positively associated with lifespan at older ages, while greater frailty showed negative associations at older ages, consistent with the expectation that healthier animals tend to survive longer. However, these relationships did not reach significance across most timepoints and groups. Most importantly, ORX240 animals did not exhibit a distinctive relationship in which greater grip strength or lower frailty consistently predicted their enhanced survival.

Together, these findings do not support the hypothesis that the longevity advantage produced by delaying castration until 240 days results from the establishment of greater muscle or physical-function reserve built during the additional period of testicular factor exposure. Although testicular factors strongly influenced body composition before castration, and early exposure may still produce lasting effects relevant to survival, the subsequent survival advantage of ORX240 was not accompanied by greater preservation of grip strength, reduced accumulation of deficits, or a stronger relationship between these functional phenotypes and lifespan. The additional survival benefit associated with testicular exposure before 240 days therefore remains unexplained by the observed differences in body composition, grip strength, or frailty, although contributions from these phenotypes cannot be fully excluded.

## Discussion

Our findings clarify when testicular factors become detrimental to male survival. Previously, it was unclear at what stage of life these factors exert their adverse effects, whether exposure during puberty or early adulthood produces lasting changes that subsequently increase mortality, or whether the elevated male mortality hazard depends on the continued presence of testicular factors. Castration at all three ages tested, including as late as 240 days, eliminated the characteristic increase in male mortality that emerged thereafter. Moreover, castration immediately after puberty at 60 days was at least as effective as prepubertal castration at 20 days, arguing against the hypothesis that the pubertal surge in testicular factors irreversibly programs the male longevity disadvantage.

Unexpectedly, delaying castration until 240 days produced the greatest survival benefit. ORX240 males lived significantly longer than ORX20 males, with significantly lower mortality hazard between 627 and 1,010 days, suggesting that earlier testicular exposure may confer lasting survival benefits. Importantly, castration at 240 days, immediately before the typical onset of male mortality, prevented the subsequent rise in mortality despite prolonged prior exposure. This suggests that excess male mortality depends on the ongoing presence of testicular factors rather than irreversible effects of earlier exposure. Together with the comparable benefits of prepubertal and postpubertal castration, these findings indicate that testicular factors have age-dependent effects on survival, potentially beneficial earlier in life but detrimental when maintained later. Although treatment cohorts were generated sequentially, survival did not differ among their corresponding littermate-matched SHAM controls (Supplementary Figure 1B), reducing the likelihood that cohort-specific differences in baseline mortality explain the greater survival of ORX240 mice. Nevertheless, smaller cohort-specific effects or differences in responses to castration cannot be entirely excluded.

The physiological basis of this apparent early-beneficial, late-detrimental transition remains unresolved. Testicular factors strongly influenced body composition throughout adulthood, with intact males maintaining greater lean mass and substantially lower fat mass than females. However, these effects were highly reversible by castration. Following castration at 240 days, lean and fat mass rapidly shifted toward the trajectories of females and males castrated earlier, indicating that adult body composition depends largely on the current hormonal environment rather than being permanently programmed by earlier testicular factor exposure. Importantly, the greater longevity of ORX240 mice was not accompanied by consistently greater grip strength or reduced frailty relative to animals castrated earlier. This argues against a simple model in which prolonged early-life androgen exposure improves later survival by permanently establishing greater muscle mass or physical-function reserve.

These findings have important translational implications. Because removal of the testes as late as 240 days was sufficient to eliminate the subsequent male-specific mortality disadvantage, interventions aimed at blocking the detrimental effects of testicular factors may not need to be initiated before puberty or early adulthood. Instead, there may be a later therapeutic window in which the harmful effects of continued testicular factor exposure can be reduced while preserving the potential benefits of earlier exposure. This raises the possibility of developing “castration mimetics”: interventions that reproduce the beneficial effects of castration without its unwanted side effects.

Our findings may also have implications for the pronounced sex differences observed in response to longevity interventions. Many interventions tested by the ITP are effective exclusively or more strongly in males^13^. One possibility is that these interventions act, directly or indirectly, by counteracting the detrimental effects of testicular factors on survival. If so, these pathways may contribute both to the higher baseline mortality of males and to their greater responsiveness to certain longevity interventions. Testing candidate longevity interventions in intact and castrated males could therefore help distinguish compounds whose benefits depend on modifying testicular factor-associated pathways from those acting through testes-independent mechanisms.

Overall, our results argue against an “earlier is better” model of castration-mediated lifespan extension^7^. Pubertal and early-adult exposure to testicular factors, including male sex hormones, does not appear to impose an irreversible mortality cost and may instead provide lasting survival benefits, whereas continued exposure later in life becomes detrimental. Identifying the physiological transition that converts testicular factors from beneficial to harmful, and the downstream pathways responsible for this shift, may improve our understanding of both the male longevity disadvantage and the marked sex specificity of many interventions that extend lifespan.

## Methods

### Animals and husbandry

Genetically heterogeneous UM-HET3 mice were generated using the breeding scheme employed by the National Institute on Aging Interventions Testing Program (ITP), by crossing BALB/cByJ *×* C57BL/6J F1 females with C3H/HeJ *×* DBA/2J F1 males, thereby maintaining comparability with historical ITP cohorts ^14^. Mice were weaned at 19–20 days of age and assigned to orchidectomy at 20, 60, or 240 days of age (ORX20, ORX60, and ORX240, respectively) or to a sham-surgery control group. An untreated female cohort was included as a female reference group. Mice were maintained according to husbandry procedures used for ITP lifespan studies^14^, with three males and five females housed per cage. Animals were inspected daily by trained personnel for morbidity and mortality. Mice meeting humane-endpoint criteria because of severe illness, rapidly progressing tumors, fighting or other conditions incompatible with continued observation were euthanized according to ITP protocol. Euthanized animals were treated as censored observations in survival analyses.

### Surgical orchidectomy

Bilateral orchidectomy was performed at 20, 60, or 240 days of age (n *≈* 60 per group) through a midline scrotal incision under inhaled isoflurane anesthesia (3–4% in 100% oxygen). Mice were positioned in dorsal recumbency, hair over the surgical site was removed using Nair depilatory cream, and the area was prepared aseptically with three alternating applications of povidone–iodine and 1 *×* PBS. The testes were exteriorized sequentially through a single midline scrotal incision. For each testis, the spermatic cord was clamped below the testis, and the testis was excised using a high-temperature cautery pen (Bovie Medical AA11), leaving the epididymis intact. After hemostasis was confirmed, the incision was closed with tissue adhesive (Vetbond). Sham-operated mice underwent the same procedure, with each testis exteriorized sequentially and returned to the scrotal sac without removal before closure. Animals recovered in a clean, warmed cage and were monitored continuously until fully ambulatory and at least daily thereafter. Deaths occurring within 14 days of surgery were assumed to be surgery related and were excluded from the data (4 in ORX240 and 2 in SHAM240).

### Lifespan assessment

Cages were checked daily for deaths, and mice were followed until spontaneous death or euthanasia at a humane endpoint according to ITP protocol. Animals were analyzed according to their assigned treatment groups, including those that died before their scheduled surgery. Natural deaths were treated as events, whereas euthanized animals were right-censored. Survival differences were assessed using the log-rank test and the Peto–Peto modification of the Gehan–Wilcoxon test, implemented using the survdiff function in R’s survival package. Pairwise P values were adjusted separately for each test using the Benjamini–Hochberg (BH) procedure. Median lifespan and 95% confidence intervals were estimated from Kaplan–Meier survival functions. Lifespan distributions were additionally visualized using individual observations and boxplots showing medians and interquartile ranges. To further characterize differences in the timing of mortality between subgroups, survival was additionally evaluated using the Temporal Efficacy Profiler (TEP) ^12^. Results are reported using the asymptotic method for confidence interval estimation, and only when the corresponding bootstrap confidence intervals overlapped with the asymptotic estimates^12^.

### Body composition

Body weight, fat mass, and lean mass were measured every 30 days using a digital scale and EchoMRI analyzer. Measurements underwent quality control for invalid dates, duplicates, implausible values, and discrepancies between body weight and summed fat and lean mass (*≥*10 g excluded; *>*2 g and *>*10% flagged for review). Longitudinal trajectories were analyzed using generalized additive mixed models (GAMMs), including group-specific cubic regression splines for age (k = 6) and mouse identity as a random effect. Models were fitted by restricted maximum likelihood, with predicted trajectories and 95% confidence intervals. Group differences in age-dependent trajectories were tested by comparing models containing a common age smooth with models containing group-specific age smooths. These comparisons were performed using maximum-likelihood fitting and likelihood-ratio chi-square tests. Pairwise trajectory comparisons used the same approach, with P values adjusted using the BH procedure. For descriptive visualization, measurements were also summarized according to predefined nominal measurement days. Inferential longitudinal analyses were based on exact chronological age rather than nominal measurement day.

### Grip strength and frailty assessment

Forelimb and total grip strength were measured longitudinally using a Chatillon Ametek grip-strength meter. Peak force was recorded across five trials, and the mean was used for analysis. Frailty was assessed using a modified 27-item clinical frailty assessment across five categories encompassing body condition, performance of simple motor tasks, and responses to sensory stimuli^15^. Each parameter was scored as 0 (no deficit), 0.5 (mild deficit), or 1 (severe deficit), and individual scores were summed to obtain a total frailty score. All three measures were analyzed using linear mixed-effects models including group, nominal timepoint, and their interaction as fixed effects, with mouse identity as a random intercept. Models were fitted by restricted maximum likelihood, with type III tests for fixed effects and BH-adjusted pairwise comparisons using estimated marginal means.

### Proportion-of-life-lived analyses

Body composition, grip strength, and frailty were additionally analyzed as a function of proportion of life lived (PLL), defined as age at measurement divided by eventual lifespan, restricting analyses to animals that died naturally. Trajectories were visualized using group-specific LOESS curves with 95% confidence intervals. Differences were assessed using GAMMs with cubic regression splines (k = 6) and mouse-specific random effects, comparing common versus group-specific PLL smooths by likelihood-ratio tests, with BH correction for pairwise comparisons.

### Associations between health measures and lifespan

Associations between forelimb grip strength, total grip strength, or frailty index and eventual lifespan were assessed using Spearman rank correlations. Analyses were restricted to animals that died naturally and therefore had known lifespan. Spearman correlations were calculated separately for each treatment group and nominal measurement timepoint. For the longitudinal correlation analyses, P values were adjusted across group-by-timepoint comparisons within each phenotype using the BH procedure.

### Statistical analysis

Statistical analyses were performed in R version 4.5.1. Longitudinal models accounted for repeated observations within animals using mouse-specific random effects. Missing longitudinal measurements were not imputed. Unless otherwise specified, statistical tests were two-sided. Statistical significance was defined as P < 0.05 for individual or omnibus tests and BH-adjusted P < 0.05 where correction for multiple comparisons was applied. Exact sample sizes differed among longitudinal analyses because of mortality and missing measurements and are reported with the corresponding results or source data. The raw data underlying these analyses will be made publicly available upon publication of the full manuscript.

## Acknowledgments

We thank Vanessa Calderon for technical assistance and training, and the staff of the Aging Animal and Functional Assessment Core of the San Antonio Nathan Shock Center for their support with animal husbandry, monitoring, and lifespan assessments.

This work was supported by the National Institutes of Health (NIH) through the National Institute on Aging (NIA) Interventions Testing Program (U01 AG022307), the San Antonio Nathan Shock Center of Excellence in Basic Biology of Aging (P30 AG013319), and grants R00 AG068239, R01 DK138035, and R01 AG084646 awarded to Shangang Zhao.

## References

[1] Steven N. Austad and Kathleen E. Fischer. Sex differences in lifespan. Cell Metabolism, 23(6): 1022–1033, 2016. doi: 10.1016/j.cmet.2016.05.019.

[2] GBD 2021 Demographics Collaborators. Global age-sex-specific mortality, life expectancy, and population estimates in 204 countries and territories and 811 subnational locations, 1950–2021, and the impact of the COVID-19 pandemic: a comprehensive demographic analysis for the Global Burden of Disease Study 2021. The Lancet, 403(10440):1989–2056, 2024. doi: 10.1016/S0140-6736(24)00476-8.

[3] Mitchell D. Wong, Anne K. Chung, W. John Boscardin, Ming Li, Hsin-ju Hsieh, Susan L. Ettner, and Martin F. Shapiro. The contribution of specific causes of death to sex differences in mortality. Public Health Reports, 121(6):746–754, 2006. doi: 10.1177/003335490612100615.

[4] Julia Francis, Barry I. Graubard, Hormuzd Katki, and Sarah S. Jackson. Sex and all-cause mortality in the US, 1999 to 2018. JAMA Network Open, 9(1):e2556299, 2026. doi: 10.1001/jamanetworkopen.2025.56299.

[5] Catherine J. Cheng, Jonathan A. L. Gelfond, Randy Strong, and James F. Nelson. Genetically heterogeneous mice exhibit a female survival advantage that is age- and site-specific: results from a large multi-site study. Aging Cell, 18(3):e12905, 2019. doi: 10.1111/acel.12905.

[6] Nisi Jiang, Catherine J. Cheng, Jonathan Gelfond, Randy Strong, Vivian Diaz, and James F. Nelson. Prepubertal castration eliminates sex differences in lifespan and growth trajectories in genetically heterogeneous mice. Aging Cell, 22(8):e13891, 2023. doi: 10.1111/acel.13891.

[7] Michael Garratt, Malgorzata Lagisz, Johanna Staerk, Christine Neyt, Michael B. Stout, José V. V. Isola, Veronica B. Cowl, Nannette Driver-Ruiz, Ashley D. Franklin, Monica M. McDonald, David M. Powell, Susan L. Walker, Jean-Michel Gaillard, Dalia A. Conde, Jean-François Lemaître, Fernando Colchero, and Shinichi Nakagawa. Sterilization and contraception increase lifespan across vertebrates. Nature, 649(8099):1264–1272, 2026. doi: 10.1038/s41586-025-09836-9.

[8] J. B. Hamilton and G. E. Mestler. Mortality and survival: comparison of eunuchs with intact men and women in a mentally retarded population. Journal of Gerontology, 24(4):395–411, 1969. doi: 10.1093/geronj/24.4.395.

[9] Kyung-Jin Min, Cheol-Koo Lee, and Han-Nam Park. The lifespan of Korean eunuchs. Current Biology, 22(18):R792–R793, 2012. doi: 10.1016/j.cub.2012.06.036.

[10] Kalynn M. Schulz, Heather A. Molenda-Figueira, and Cheryl L. Sisk. Back to the future: the organizational-activational hypothesis adapted to puberty and adolescence. Hormones and Behavior, 55(5):597–604, 2009. doi: 10.1016/j.yhbeh.2009.03.010.

[11] Nisi Jiang, Jonathan Gelfond, Qianqian Liu, Randy Strong, and James F. Nelson. The Gehan test identifies life-extending compounds overlooked by the log-rank test in the NIA Interventions Testing Program: Metformin, Enalapril, caffeic acid phenethyl ester, green tea extract, and 17-dimethylaminoethylamino-17-demethoxygeldanamycin hydrochloride. GeroScience, 46(5): 4533–4541, 2024. doi: 10.1007/s11357-024-01161-9.

[12] Nisi Jiang, Catherine J. Cheng, Qianqian Liu, Randy Strong, Jonathan Gelfond, and James F. Nelson. Deciphering the timing and impact of life-extending interventions: temporal efficacy profiler distinguishes early, midlife, and senescence phase efficacies. Nature Communications, 16:10164, 2025. doi: 10.1038/s41467-025-65158-4.

[13] Nisi Jiang, Ziying Xu, Shangang Zhao, Jonathan Gelfond, Randy Strong, and James F. Nelson. Sex as a major determinant of pro-longevity drug efficacy: a review of two decades of the NIA Interventions Testing Program. The Journals of Gerontology: Series A, 80(8):glaf138, 2025. doi: 10.1093/gerona/glaf138.

[14] Richard A. Miller, David E. Harrison, Clinton M. Astle, Robert A. Floyd, Kevin Flurkey, Kenneth L. Hensley, Martin A. Javors, Christiaan Leeuwenburgh, James F. Nelson, Ennio Ongini, Nancy L. Nadon, Huber R. Warner, and Randy Strong. An Aging Interventions Testing Program: study design and interim report. Aging Cell, 6(4):565–575, 2007. doi: 10.1111/j.1474-9726.2007.00311.x.

[15] J. C. Whitehead, B. A. Hildebrand, M. Sun, M. R. Rockwood, R. A. Rose, K. Rockwood, and S. E. Howlett. A clinical frailty index in aging mice: comparisons with frailty index data in humans. The Journals of Gerontology: Series A, Biological Sciences and Medical Sciences, 69 (6):621–632, 2014. doi: 10.1093/gerona/glt136.

