## Supplementary Information for "Beneficial early, harmful later: age-dependent effects of testicular factors on male mortality"

### Supplementary Figures

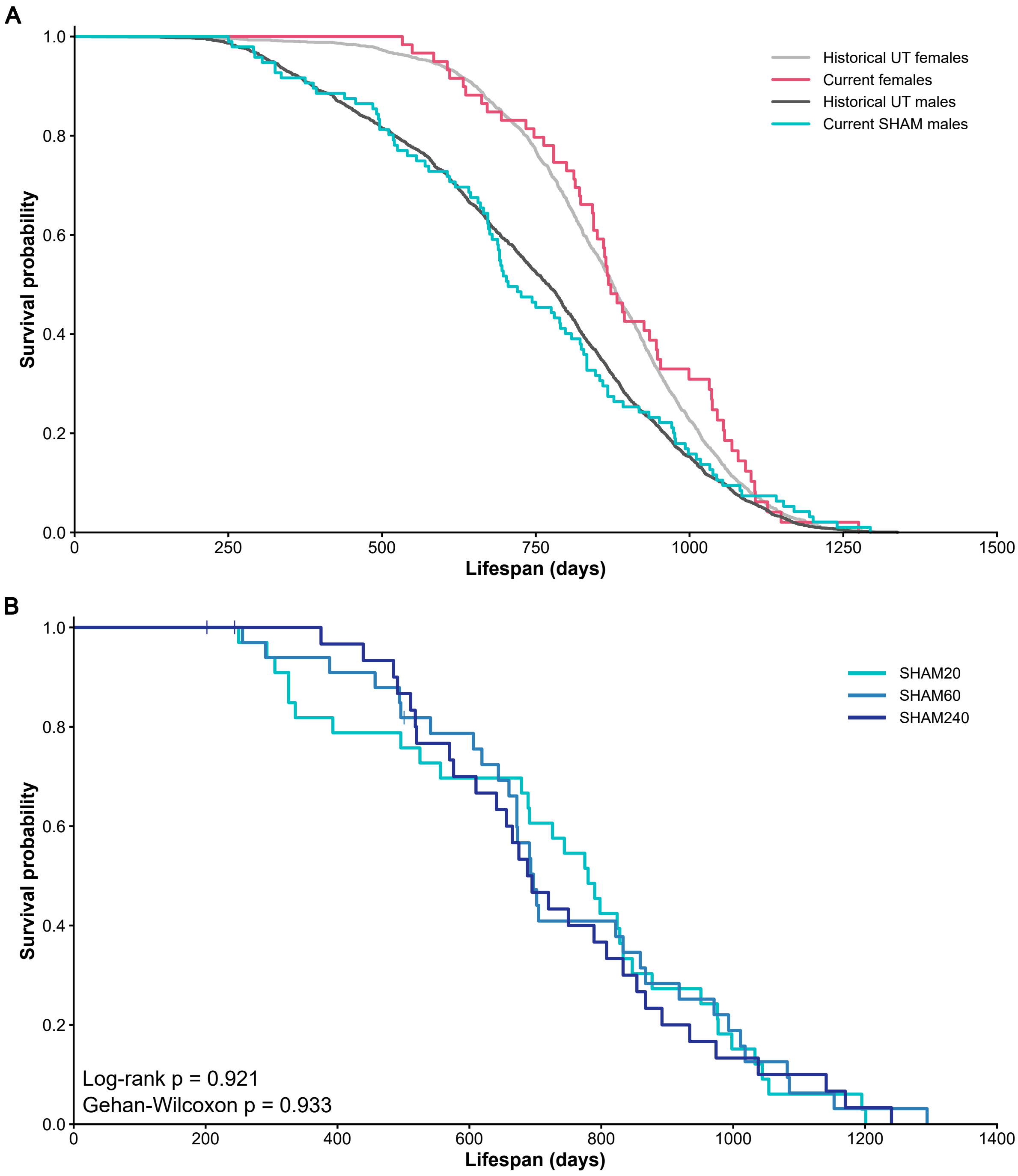


**Supplementary Figure 1. Survival of current study controls compared with historical UM-HET3 cohorts and survival across SHAM surgery groups.** (A) Kaplan–Meier survival curves comparing females and pooled SHAM males from the current study with historical female and male UM-HET3 controls from the University of Texas site of the NIA Interventions Testing Program. The current SHAM cohort had a lower median lifespan than the historical male cohort, but the overall survival distributions did not differ significantly. (B) Kaplan–Meier survival curves for males receiving sham surgery at 20, 60, or 240 days of age (SHAM20, n=33; SHAM60, n=33; SHAM240, n=33). Survival did not differ significantly among sham-surgery cohorts (log-rank, P = 0.921; Gehan–Wilcoxon, P = 0.933), supporting their pooling into a single SHAM male group for primary analyses.

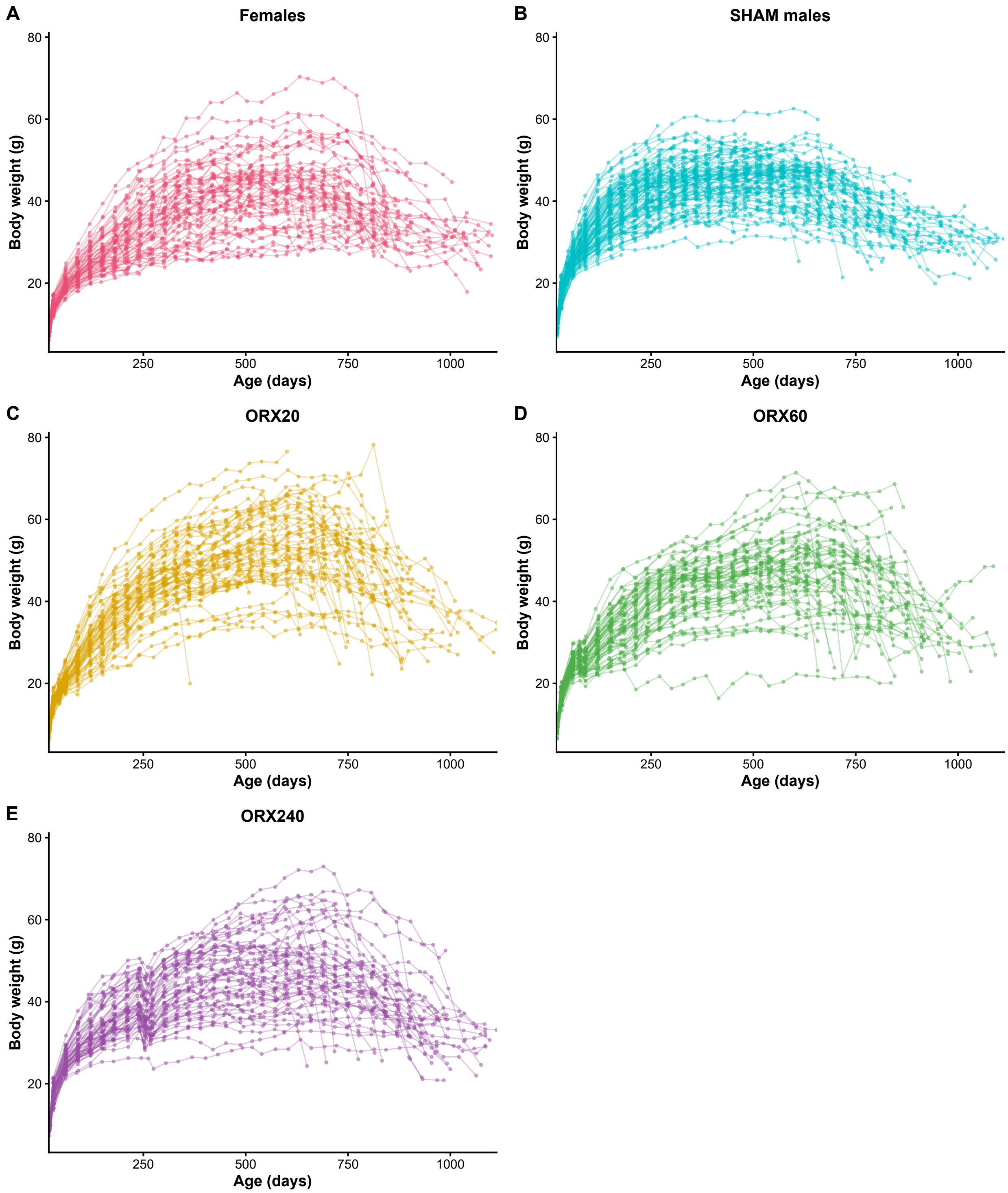


**Supplementary Figure 2. Individual longitudinal body-weight trajectories across treatment groups.** Individual body-weight trajectories for (A) females, (B) SHAM males, (C) ORX20, (D) ORX60, and (E) ORX240 mice across chronological age. Connected points represent repeated measurements from individual animals. These plots illustrate within-animal variation underlying the group-level body-weight trajectories presented in Figure 2.

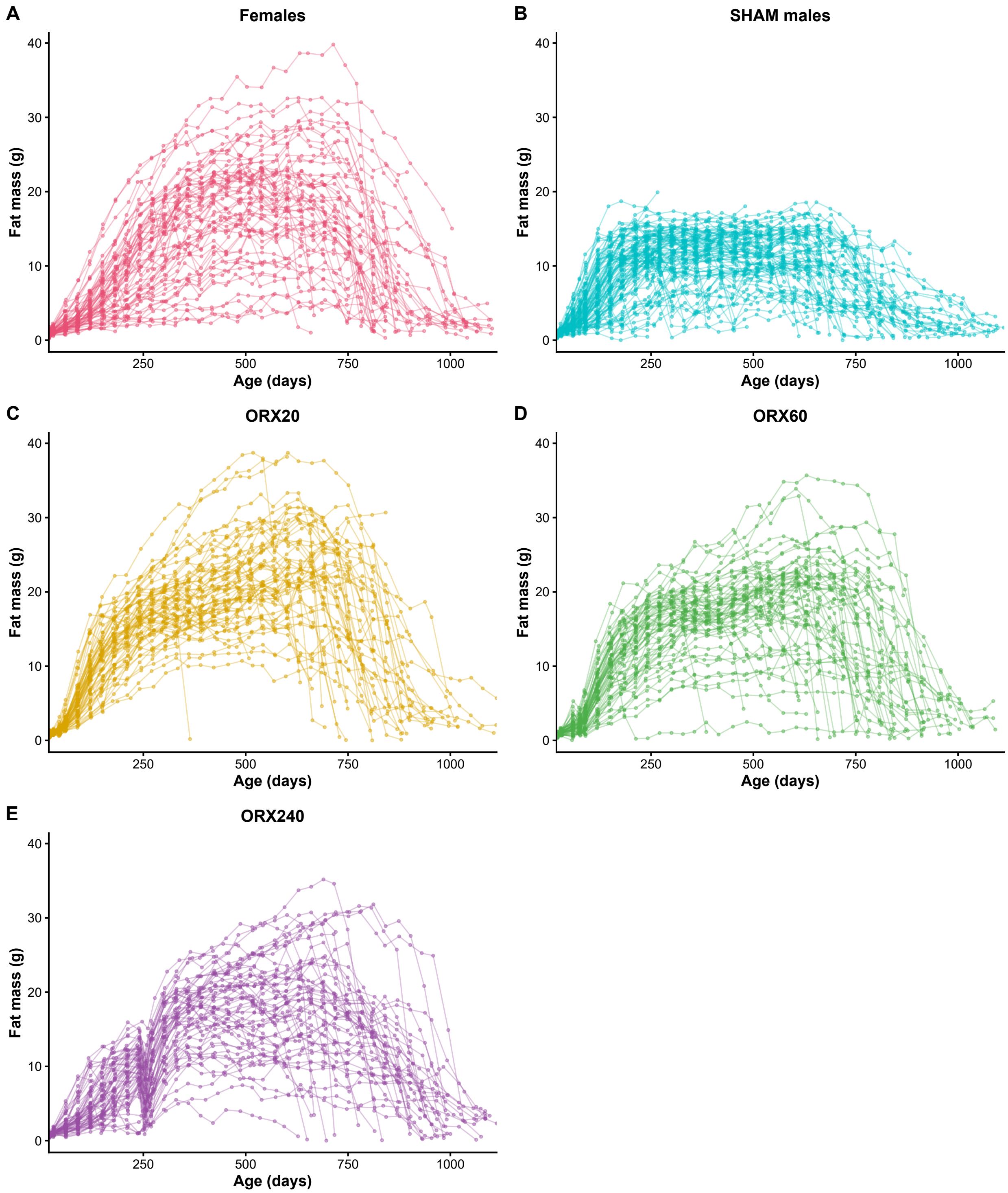


**Supplementary Figure 3. Individual longitudinal fat-mass trajectories across treatment groups.** Individual fat-mass trajectories for (A) females, (B) SHAM males, (C) ORX20, (D) ORX60, and (E) ORX240 mice across chronological age. Connected points represent repeated measurements from individual animals. Individual trajectories demonstrate the substantially lower adiposity of intact SHAM males and the increased fat accumulation following castration at each age.

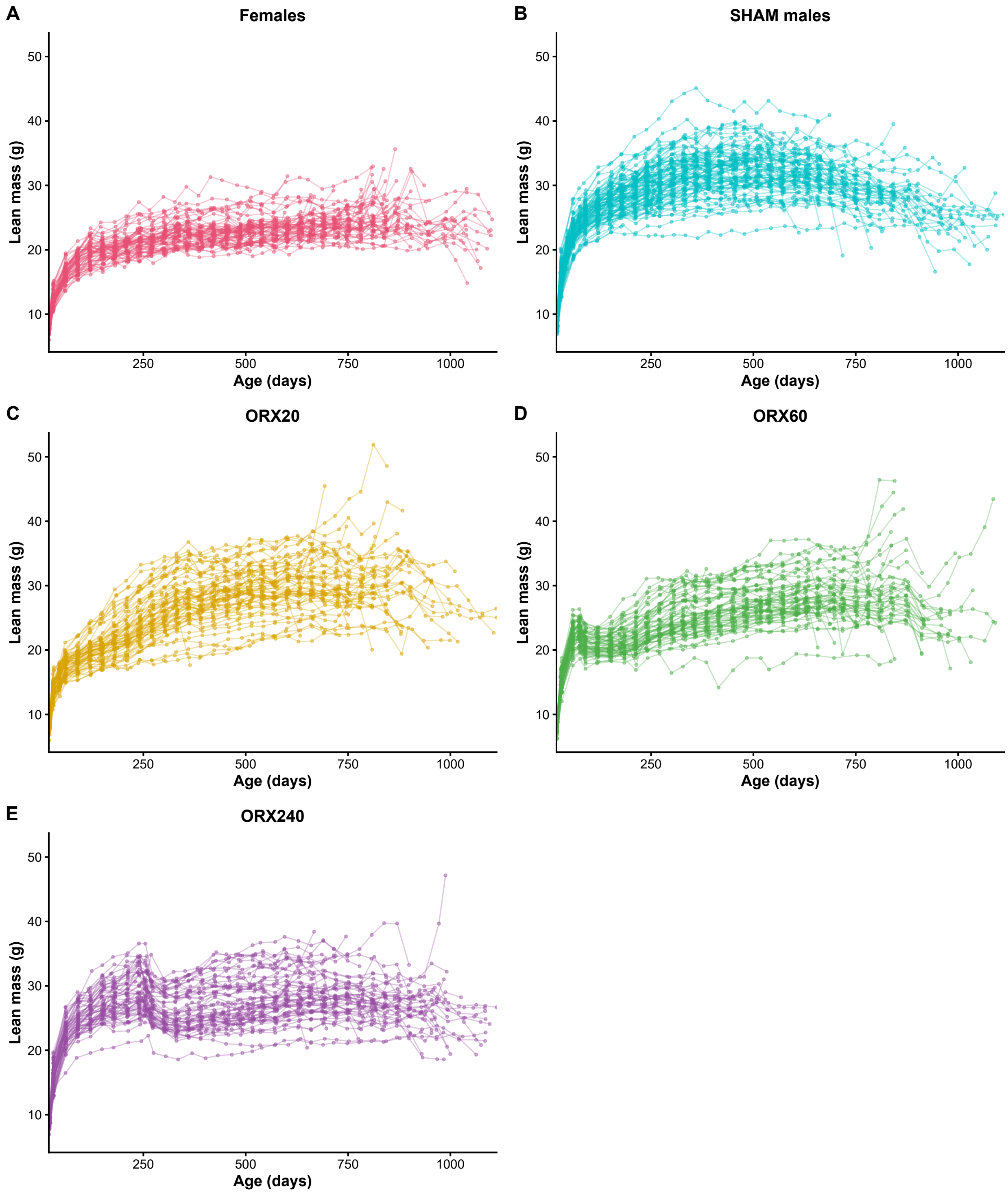


**Supplementary Figure 4. Individual longitudinal lean-mass trajectories across treatment groups.** Individual lean-mass trajectories for (A) females, (B) SHAM males, (C) ORX20, (D) ORX60, and (E) ORX240 mice across chronological age. Connected points represent repeated measurements from individual animals. Individual trajectories illustrate the greater lean mass of intact males and the shift toward lower lean mass following castration.

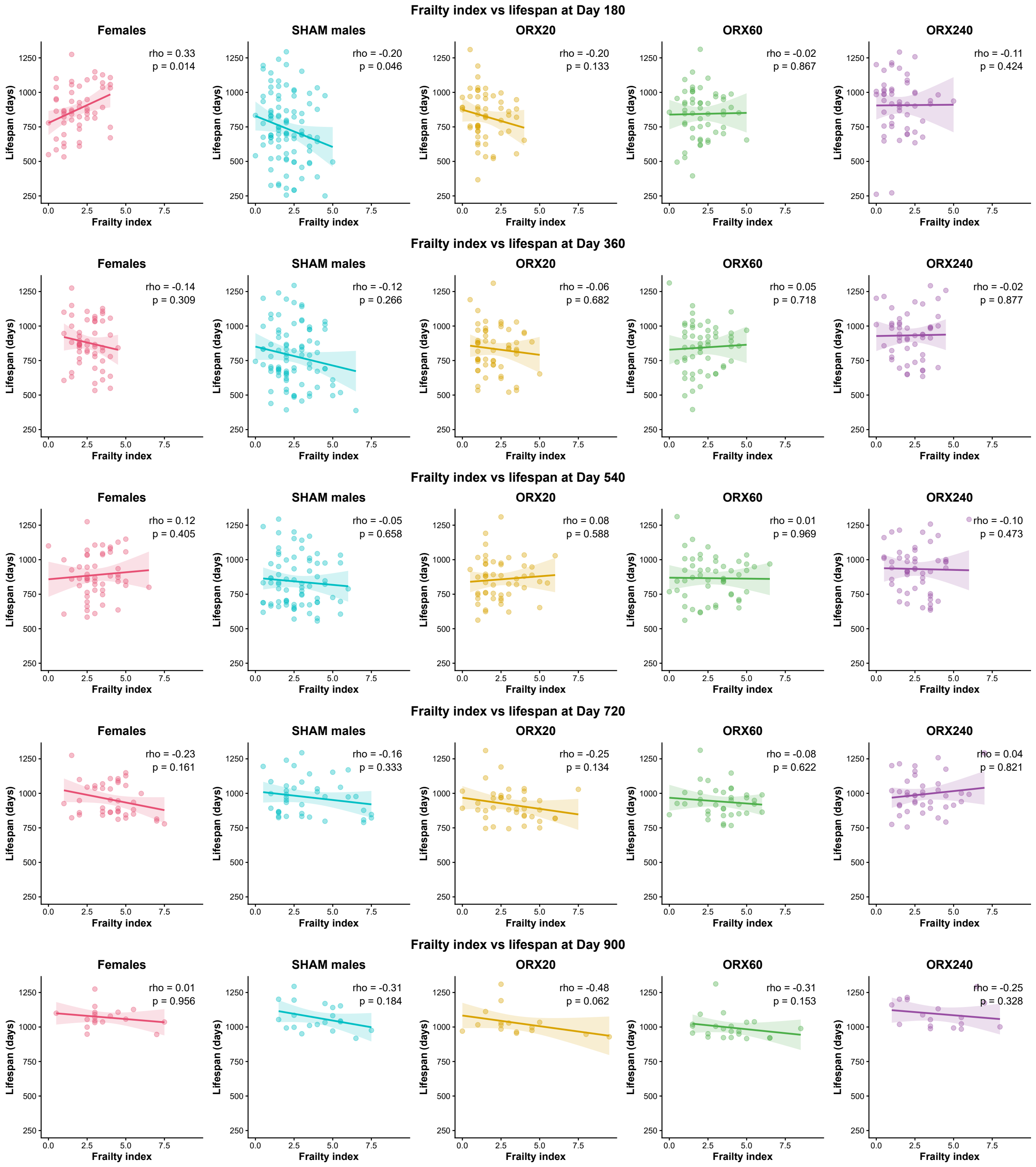


**Supplementary Figure 5. Associations between frailty and lifespan across age and treatment groups.** Spearman rank correlations between frailty index and eventual lifespan in females, SHAM males, ORX20, ORX60, and ORX240 mice at 180, 360, 540, 720, and 900 days of age. Points represent individual animals, lines show the fitted relationship for visualization, and each panel reports the Spearman correlation coefficient ($\rho$) and corresponding P value. Analyses were restricted to animals that died naturally and therefore had known lifespans. Associations between frailty and lifespan varied across ages and treatment groups, with no consistent relationship uniquely explaining the greater longevity of ORX240 mice. This provides a more detailed presentation of the correlations summarized in the rightmost panel of Figure 3C.
